# Evaluation of Mosaicism and Off Target Mutations in CRISPR-Mediated Genome Edited Bovine Embryos

**DOI:** 10.1101/2020.06.04.134759

**Authors:** Sadie L. Hennig, Joseph R. Owen, Jason C. Lin, Amy E. Young, Pablo J. Ross, Alison L. Van Eenennaam, James D. Murray

**Author notes:** Sadie L. Hennig and Joseph R. Owen contributed equally.

## Abstract

The CRISPR/Cas9 genome editing tool has the potential to improve the livestock breeding industry by allowing for the introduction of desirable traits. Although an efficient and targeted tool, the CRISPR/Cas9 system can have some drawbacks, including off-target mutations and mosaicism, particularly when used in developing embryos. Here, we introduced genome editing reagents into single-cell bovine embryos to compare the effect of Cas9 mRNA and protein on the mutation efficiency, level of mosaicism, and evaluate potential off-target mutations utilizing next generation sequencing. We designed guide-RNAs targeting three loci (POLLED, H11, and ZFX) in the bovine genome and saw a significantly higher rate of mutation in embryos injected with Cas9 protein (84.2%) vs. Cas9 mRNA (68.5%). In addition, the level of mosaicism was higher in embryos injected with Cas9 mRNA (100%) compared to those injected with Cas9 protein (94.2%), with little to no unintended off-target mutations detected. This study demonstrates that the use of Cas9 protein, rather than Cas9 mRNA, results in a higher editing efficiency in bovine embryos while lowering the level of mosaicism. However, further optimization must be carried out for the CRISPR/Cas9 system to become feasible for single-step embryo editing in a commercial system.

## INTRODUCTION

CRISPR-mediated genome editing in livestock zygotes offers an attractive approach to introduce useful genetic variation into the next generation of cattle breeding programs. However, genetic mosaicism is particularly problematic for CRISPR-mediated genome editing in developing zygotes^1,2^. Genetic mosaicism complicates phenotypic analysis of F0 animals and may complicate screening multiple founders and breeding mosaic founders to produce an F1 generation. While this is routine in plant and mouse research, such approaches are time-consuming and essentially cost-prohibitive in large food animal species with long generation intervals like cattle.

A limited number of genome editing studies have been reported in bovine zygotes^3^, and indicate the frequent production of mosaic embryos. The frequency of mosaicism varies depending upon the type of site-directed nuclease used, the timing of editing relative to embryonic development, the form and efficiency of the targeting regents, the intrinsic properties of the target locus, and the method of delivery^1^.

Correspondingly, there are a number of experimental variables that need to be optimized to improve the efficiency of obtaining non-mosaic, homozygous genome edited founder cattle. In this study, we focused on the type of CRISPR/Cas9 system delivered (i.e. mRNA or protein) and report the impact on mutation efficiency, levels of mosaicism, and off-target mutations based on next generation sequencing when using CRISPR-mediated genome editing of bovine zygotes.

## RESULTS

### Guide construction and testing

To determine the optimal parameters for CRISPR/Cas9-mediated genome editing in bovine zygotes, efficiency following microinjection was investigated for three gRNA per locus on three different chromosomes. Three gRNAs were designed targeting the POLLED locus on chromosome 1, a safe harbor locus (H11) on chromosome 17 and a locus (ZFX) on the X-chromosome downstream of the Zinc Finger, X-linked gene (Supplementary Information, Table S1). Three gRNAs per locus were independently injected alongside Cas9 protein in groups of 30 zygotes, 18 hours post insemination (hpi). Groups of 50 non-injected embryos were cultured as controls. The highest mutation rates were 76.9% for gRNA2 targeting the POLLED locus, 83.3% for gRNA1 targeting the H11 locus, and 77.8% for gRNA3 targeting the ZFX locus (Supplementary Information, Table S2; χ^2^ test, P < 0.05). Overall, there was a decrease in the number of embryos that reached the blastocyst stage as the rate of mutation for a given gRNA increased. For each locus, the gRNA with the highest mutation rate was associated with the lowest developmental rate (Supplementary Information, Table S2). gRNAs with the highest mutation rate were selected for further analysis.

Guides targeting the POLLED locus, the H11 locus and the ZFX locus were then injected in groups of 30 *in vitro* fertilized embryos 18hpi alongside either Cas9 mRNA or protein (Table 1). While there was no significant difference in development to the blastocyst stage when comparing embryos injected with Cas9 mRNA (16.2%) or Cas9 protein (16.4%), there was a significant decrease in the proportion of zygotes reaching the blastocyst stage for both groups compared to non-injected controls (30.7%; Fig. 1a; χ^2^ test, P < 0.05). Mutation rates were significantly higher for Cas9 protein (84.2%) compared to Cas9 mRNA (68.5%) (Fig. 1b; χ^2^ test, P < 0.05) for all three loci located on different chromosomes (Fig. 1c).

**Table 1.** Number of zygotes reaching the blastocyst developmental stage following microinjection of either Cas9 mRNA or protein and gRNAs targeting three loci (POLLED, H11, and ZFX) on different chromosomes. I*n vitro* fertilized bovine embryos were injected 18 hours post insemination, and the percentage of blastocysts with Cas9-induced mutations was determined by sequence analysis. Letters that differ in the same column are significantly different (P < 0.05).

| Cas9 | gRNA | Injected Groups | Total Embryos | Total Blasts (%) | Total Analyzed | Total Mutation (%) |
| --- | --- | --- | --- | --- | --- | --- |
| mRNA | control | - | 492 | 131 (27)<sup>a</sup> | - | - |
|  | POLLED | 4 | 109 | 21 (19)<sup>b</sup> | 22 | 16 (73)<sup>a</sup> |
|  | H11 | 7 | 191 | 28 (15)<sup>b</sup> | 27 | 19 (70)<sup>a</sup> |
|  | ZFX | 14 | 372 | 60 (16)<sup>b</sup> | 62 | 41 (67)<sup>a</sup> |
| protein | control | - | 749 | 250 (33)<sup>a</sup> | - | - |
|  | POLLED | 12 | 316 | 53 (17)<sup>b</sup> | 42 | 36 (86)<sup>b</sup> |
|  | H11 | 6 | 162 | 27 (17)<sup>b</sup> | 39 | 35 (90)<sup>b</sup> |
|  | ZFX | 22 | 562 | 91 (16)<sup>b</sup> | 90 | 73 (81)<sup>b</sup> |

**Figure 1.**
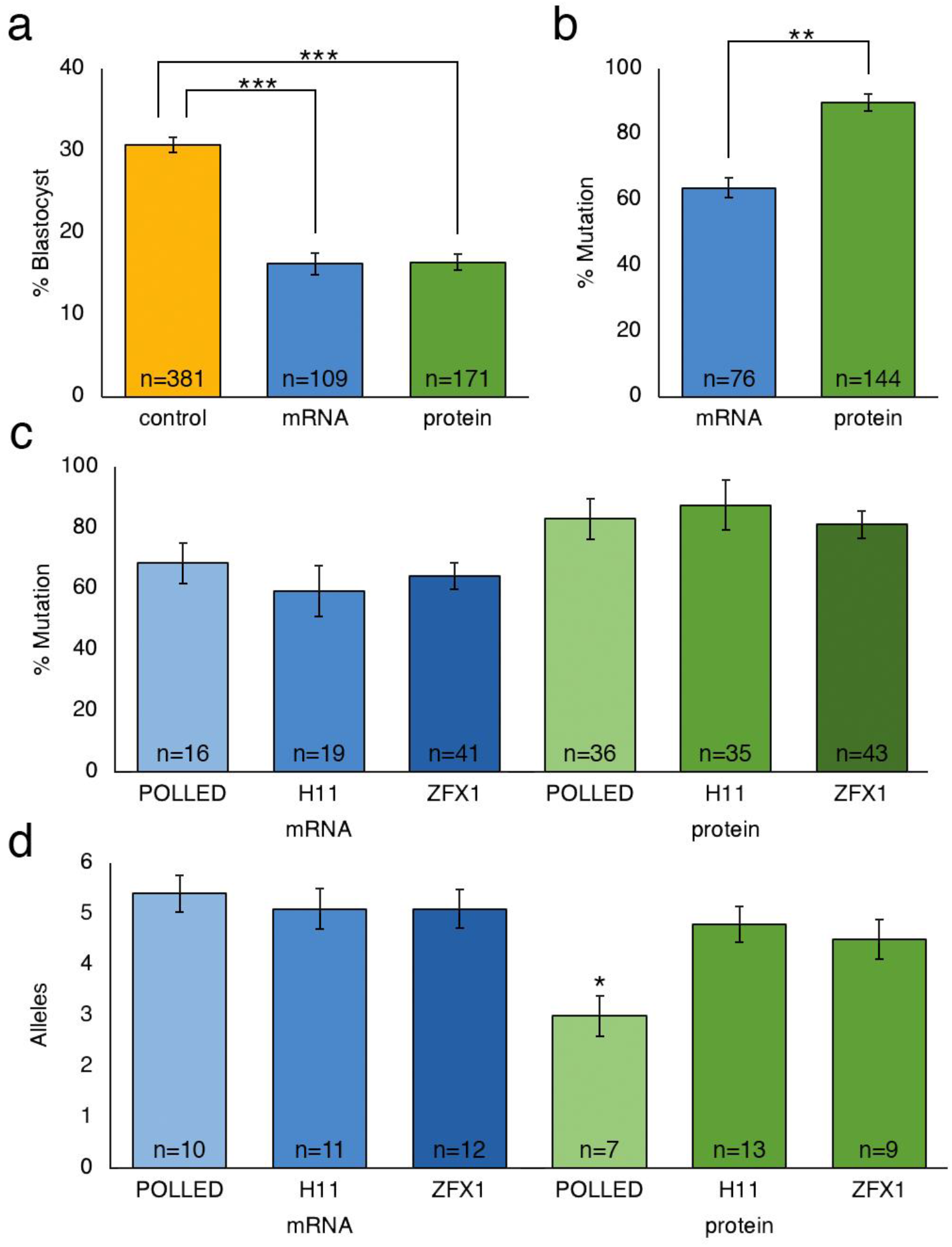
Percentage of uninjected control and microinjected zygotes reaching the blastocyst developmental stage following microinjection of either Cas9 mRNA or protein into *in vitro* fertilized bovine embryos 18 hours post insemination, and percentage analyzed blastocysts with Cas9-induced mutations. (a) Blastocyst developmental percentage of CRISPR injected zygotes for all three loci compared to control non-injected zygotes. (b) Percentage of blastocysts with Cas9-induced mutations when injecting either Cas9 mRNA or protein alongside gRNAs targeting all three loci. (c) Percentage of blastocysts with Cas9 mRNA or protein-induced mutation by and gRNAs targeting three loci (POLLED, H11, and ZFX) in the bovine genome. Error bars = standard error of the mean. (d) Average number of alleles per blastocyst when injecting Cas9 mRNA or protein targeting three loci (POLLED, H11, and ZFX) in the bovine genome. *P < 0.05 **P < 0.005 ***P < 0.0005.

### Evaluation of mosaicism and off-target insertions and deletions

To evaluate the level of mosaicism, 69 blastocysts (19 gRNA2 targeting the POLLED locus (10 Cas9 mRNA, 9 Cas9 protein), 26 gRNA1 targeting the H11 locus (11 Cas9 mRNA, 15 Cas9 protein), and 24 targeting the ZFX locus (13 Cas9 mRNA, 11 Cas9 protein)) were collected, barcoded by PCR amplification and sequenced on a PacBio sequencer (Supplementary Information, Table S3). Consensus sequences were called from raw reads using circular consensus sequencing (ccs) with a minimum of 3 passes, a minimum predicted accuracy of 99% and a maximum length of 700bp (Supplementary Information, Table S4). Unsorted ccs reads were aligned to each of the target sequences to analyze the types of insertions/deletions (indels) surrounding the predicted cut site with 26,460 reads aligned to the POLLED target site; 78,305 reads aligned to the H11 target site; and 66,780 reads aligned to ZFX target site (Supplementary Information, Table S5). About half of the aligned sequences for the POLLED locus were wild type sequences (47.8%), while almost three quarters of the H11 and ZFX reads were wild type sequences (75.7% and 71.3%, respectively). The primary indels for reads aligned to the POLLED locus were 7bp deletion (1672 reads), 11bp deletion (1751), 4bp deletion (6356 reads) and 1bp insertion (2250 reads); aligned to the H11 locus were 11bp deletion (3246 reads), 6bp deletion (3813 reads), 3bp deletion (4091 reads), and 1bp deletion (7853 reads); and aligned to the ZFX locus were 14bp deletion (4222 reads), 9bp deletion (2998 reads), 3bp deletion (3198 reads), 1bp deletion (2194 reads) and 1bp insertion (6532 reads) (Supplementary Information, Table S5).

Ccs reads were then sorted by barcode and analyzed by individual embryos (Fig. 2). Seven samples were discarded from further analysis due to a lack of reads following the quality filtering step (Supplemental Table S3). A total of 10 samples contained only wild type sequence (7 Cas9 mRNA and 3 Cas9 protein), resulting in an overall mutation rate of ~84% (Table 2). Of the 62 samples injected 18hpi, four contained only mutated alleles, without evidence for any wild type sequence. All four samples were from embryos injected with Cas9 protein (Supplementary Information, Table S6). Three of these samples contained only one allele and were presumably non-mosaic homozygous, although our analyses could not rule out an unmappable mutation (e.g. large insertion) at the second allele. Each of the mutated embryos containing more than a single allele had at least three individual alleles or a disproportion of reads for each allele, for example 75% wildtype and 25% mutant (Supplementary Information, Figure S1), suggesting these embryos were mosaic rather than heterozygous. This translates to 94.2% mosaicism when injecting Cas9 protein compared to 100% mosaicism when injecting Cas9 mRNA.

**Table 2.** Editing efficiencies, mosaicism, average number of alleles and percent wild type reads as determined by PacBio sequencing of 63 blastocysts following microinjection of Cas9 mRNA or protein alongside gRNAs targeting three loci (POLLED, H11, and ZFX) on different chromosomes. I*n vitro* fertilized bovine embryos were injected 18 hours post insemination. Letters that differ in the same column are significantly different (P < 0.05). SEM = standard error of the mean.

| Locus | n | Cas9 | % non-edited | % edited<br>non-mosaic | % mosaic<br>embryos | Alleles | SEM | % Wild Type | SEM |
| --- | --- | --- | --- | --- | --- | --- | --- | --- | --- |
| POLLED | 10 | mRNA | 0.0 | 0.0 | 100.0 | 5.4<sup>a</sup> | ±0.365 | 42.5<sup>a</sup> | ±7.52 |
|  | 7 | protein | 0.0 | 14.3 | 85.7 | 3.0<sup>b</sup> | ±0.398 | 9.1<sup>b</sup> | ±8.11 |
| H11 | 11 | mRNA | 36.4 | 0.0 | 100.0 | 5.1<sup>a</sup> | ±0.396 | 70.9<sup>a</sup> | ±7.01 |
|  | 13 | protein | 15.4 | 7.7 | 92.3 | 4.8<sup>a</sup> | ±0.353 | 33.7<sup>b</sup> | ±6.69 |
| ZFX | 12 | mRNA | 25.0 | 0 | 100.0 | 5.1<sup>a</sup> | ±0.375 | 79.7<sup>a</sup> | ±6.94 |
|  | 9 | protein | 11.1 | 11.1 | 88.9 | 4.5<sup>a</sup> | ±0.386 | 43.5<sup>b</sup> | ±7.47 |

**Figure 2.**
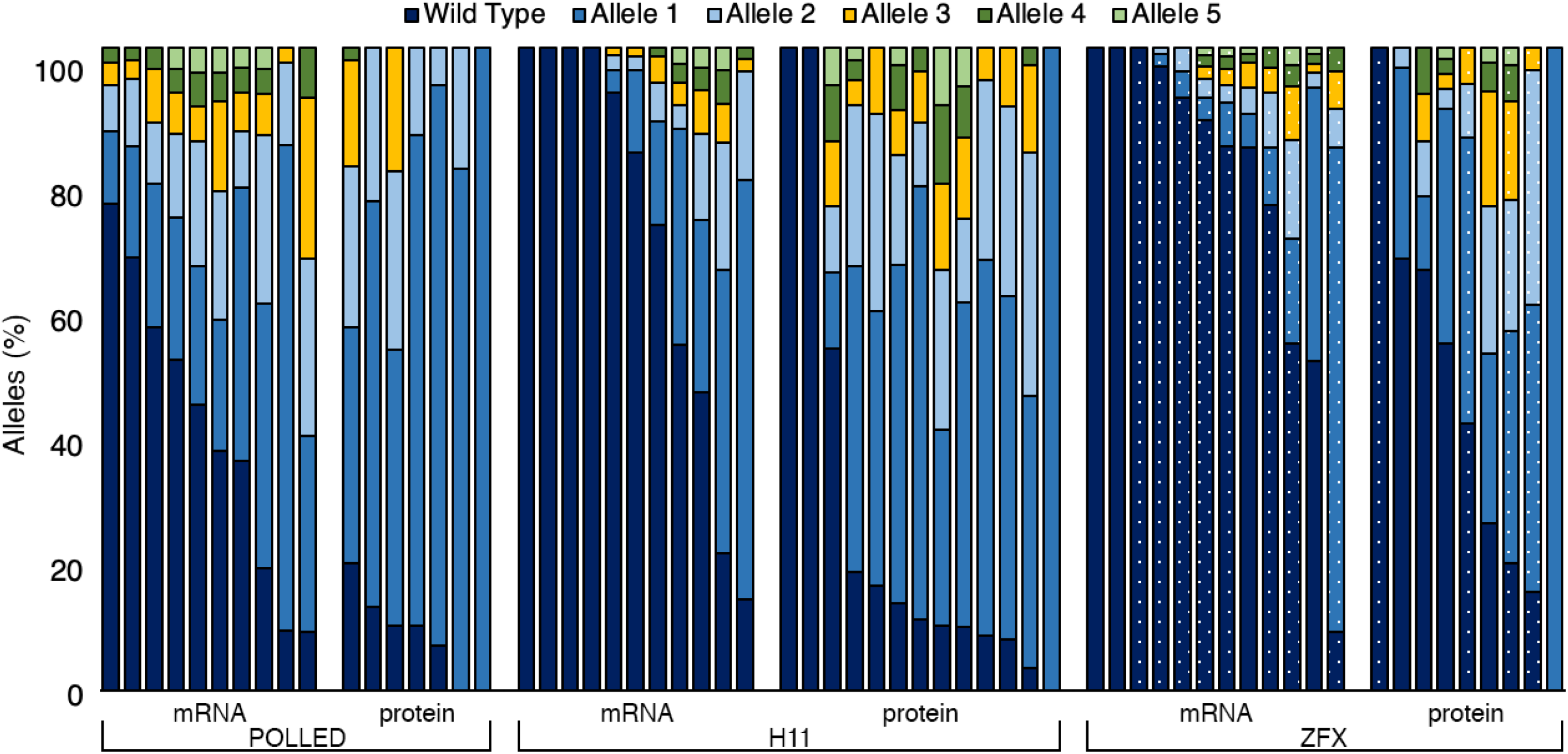
Bar graph depicting the percentage of alleles determined by PacBio sequencing in each of the 62 blastocysts microinjected 18 hours post insemination with either Cas9 mRNA or protein and gRNAs targeting the POLLED, H11 and ZFX loci. For ZFX locus: dotted bars are female; solid bars are male.

There was a decreased average number of alleles (3.0 ± 0.4) when targeting the POLLED locus using Cas9 protein (Fig 1d; Table 2). There was no significant difference in the number of alleles for the other loci when injecting Cas9 mRNA or protein. However, there was a significant increase in the number of alleles when comparing polled samples of embryos injected 18hpi with guides alongside Cas9 mRNA (5.23± 0.268), as compared to protein (4.23 ± 0.268) (ANOVA, P < 0.05). In addition, there was a significant increase in the percentage of wild type alleles present when injecting Cas9 mRNA compared to Cas9 protein for each of the three loci (42.5% vs. 9.1%, 70.9% vs. 33.7% and 79.7% vs. 43.5%, respectively; P < 0.05).

A total of 24 potential off-target sites were predicted across 11 bovine chromosomes (1, 4, 7, 8, 10, 12, 14, 18, 21, 27 and X) (Supplementary Information, Table S7) for the three loci. The 24 predicted off-target sites were PCR amplified, barcoded and sequenced using an Illumina MiSeq sequencer for each of the 69 samples (Supplementary Information, Table S3). HTStream processed reads were aligned to the 24 predicted sites with 10,399,614 reads mapped with coverage ranging from 1X to 112X per sample per site (Supplementary Information, Table S7). Genetic variation was found throughout the samples in each of the 24 predicted off-target sites with almost no indels present at the predicted off-target cut site with the exception of two targets. A 12bp deletion 26bp downstream from a predicted off-target cut site for the H11 gRNA targeting chr1: 7454978 was detected in 69,434 reads (6.8%) (Supplemental Information, Table S7). Additionally, 2,397 reads (0.51%) contained a 3bp deletion 11bp downstream from the predicted off-target cut site of the ZFX gRNA target chr21: 28506796 (Supplemental Information, Table S7).

## DISCUSSION

The ability to efficiently generate non-mosaic, homozygous founder animals is important for the production of genome edited livestock. The use of the CRISPR/Cas9 system has been reported across many livestock species^3^, but few reports have characterized its use in bovine embryos. In this study, using the CRISPR/Cas9 system, we identified gRNAs that resulted in high rates of mutation at target locations in two autosomes and the X chromosome in bovine embryos with an overall high efficiency (81-90%). Significant differences were observed in gRNA efficiency within a locus, but not between loci. Microinjection of CRISPR/Cas9 editing reagents in zygotes reduced development to the blastocyst stage compared to non-injected controls. However, no difference was observed in the number of embryos that reached the blastocyst stage when comparing embryos injected with Cas9 mRNA or protein (16.2% vs. 16.4%). This finding was important because we observed a significantly higher rate of mutation in blastocysts when injecting Cas9 protein compared to Cas9 mRNA (84.2% and 68.5%, respectively). This difference is likely due to the immediate availability of the gRNA/Cas9 ribonucleoprotein (RNP) complex to induce mutation in the embryo. When Cas9 mRNA is injected, there is a delay in genome editing as Cas9 mRNA must be translated into protein before it can combine with the gRNA to induce a DSB^4^.

Mosaicism, the presence of more than two alleles in an individual, is a common problem in livestock genome editing^5^, with a high rate of embryos resulting in multiple alleles (Table 3). Studies utilizing transcription activator-like effector nucleases (TALENs) have demonstrated lower mosaicism rates than we observed here; however, the proportion of edited embryos tends to be lower as well^6,7^. A study employing a zinc finger nuclease (ZFN) in bovine embryos demonstrated both high embryo editing efficiency and mosaicism rates as compared to those found in TALEN edited embryos^8^. However, the prevalence of mosaicism was reduced when injecting embryos at 8hpi compared to 18hpi, before S-phase had occurred^8^. While we were able to induce mutations in embryos at a high rate, we also observed a high level of mosaicism when injecting 18hpi. Many studies of editing in livestock zygotes similarly report high levels of mosaicism when utilizing CRISPR/Cas9 (Table 3). Many of these studies characterized mosaicism by sequencing the PCR amplicon of the genomic regions flanking the gRNA target sequence and then decomposing the resulting chromatogram data with the TIDE bioinformatics package^9^. Although this approach is cost-effective and rapid, next generation sequencing of the PCR products allows for a more accurate characterization of the different alleles that are present in a mosaic individual, and their relative abundance^10^.

**Table 3.** Published results of genome editing targeting the NHEJ pathway in livestock zygotes, and rates of mosaicism (where available). Modified from Mclean et al.^3^. ^a^Transcription activator-like effector (TALE), zinc finger (ZF). ^b^Nuclease delivered as plasmid, mRNA, or ribonucleoprotein (RNP) complex. ^c^Cytoplasmic injection (CI) or electroporation (E). ^d^*In vitro* fertilized (IVF) or parthenogenetic (PG) embryos. ^e^ normalized on the total number of edited embryos or not determined (ND).

| Nuclease<sup>a</sup> | Reagent<sup>b</sup> | Animal | Delivery Method<sup>c</sup> | Delivery time (post IVF)/h<sup>d</sup> | Target locus | Edited embryos % | Mosaic embryos %<sup>e</sup> | Edited offspring | Mosaic offspring | Reference |
| --- | --- | --- | --- | --- | --- | --- | --- | --- | --- | --- |
| TALE | mRNA | Bovine | CI | 19 | *ACAN* or *GDF8* | 52 | 20 | - | - | 6 |
| TALE | mRNA | Bovine | CI | 24 | *GDF8* | 31-67 | ND | 3/4 | 1/3 | 7 |
| TALE | mRNA | Ovine | CI | 24 | *GDF8* | ND | ND | 1/9 | 0/1 | 7 |
| ZF | Plasmid | Bovine | CI | 8 | *LGB* | 71 | 100 | - | - | 8 |
| ZF | Plasmid | Bovine | CI | 18 | *LGB* | 83 | 100 | - | - | 8 |
| ZF | mRNA | Bovine | CI | 8 | *LGB* | 70 | 75 | - | - | 8 |
| Cas9 | Plasmid | Porcine | CI | 17 | *GGTA1* | ND | ND | 11/12 | 4/11 | 42 |
| Cas9 | mRNA | Ovine | CI | 0 | *PDX1* | 67 | 38 | 2/4 | 2/2 | 10 |
| Cas9 | mRNA | Ovine | CI | 6 | *PDX1* | 60 | 67 | - | - | 10 |
| Cas9 | mRNA | Ovine | CI | 14 | *BMPR-IB* | 38 | 86 | - | - | 43 |
| Cas9 | mRNA | Ovine | CI | 22 | *MSTN* | 50 | 80 | 10/22 | 4/10 | 44 |
| Cas9 | mRNA | Porcine | CI | 3 | *Tet1* | 94 | 30 | - | - | 45 |
| Cas9 | mRNA | Porcine | CI | 8 | *Tet1* | 100 | 33 | - | - | 45 |
| Cas9 | mRNA | Porcine | CI | 18 | *Tet1* | 83 | 100 | - | - | 45 |
| Cas9 | mRNA | Porcine | CI | 17 | *Npc1l1* | 88 | ND | 11/11 | 9/11 | 46 |
| Cas9 | RNP | Bovine | CI | 10 (IVF),<br>1 (PG) | *POU5F1* | 86 | 34 | - | - | 47 |
| Cas9 | RNP | Bovine | E | 10 | *GDF8* | 27-67 | 75-100 | - | - | 48 |
| Cas9 | RNP | Bovine | E | 15 | *GDF8* | 19-67 | 92-100 | - | - | 48 |
| Cas9 | RNP | Porcine | CI | 0 | *GalT* | 21 | 100 | - | - | 49 |
| Cas9 | RNP | Porcine | CI | 0 + 6 | *GalT* | 23 | 100 | - | - | 49 |
| Cas9 | RNP | Porcine | CI | 6 | *GalT* | 65 | 82 | - | - | 49 |
| Cas9 | RNP | Porcine | E | 12 | *TP53* | 88 | 52 | 6/9 | 5/6 | 50 |
| Cas9 | mRNA | Bovine | CI | 0 | *PAEP* or<br>*CSN2* | 88 | 30 | - | - | 12 |
| Cas9 | RNP | Bovine | CI | 0 |  | 87 | 30 | - | - | 12 |
| Cas9 | RNP | Bovine | CI | 10 |  | 83 | 35 | - | - | 12 |
| Cas9 | mRNA | Bovine | CI | 20 |  | 84 | 100 | - | - | 12 |
| Cas9 | RNP | Bovine | CI | 20 |  | 83 | 100 | - | - | 12 |
| Cas9 | mRNA | Bovine | CI | 18 | POLLED | 73 | 100 | - | - | This study |
| Cas9 | RNP | Bovine | CI | 18 | POLLED | 86 | 86 | - | - | This study |
| Cas9 | mRNA | Bovine | CI | 18 | H11 | 70 | 100 | - | - | This study |
| Cas9 | RNP | Bovine | CI | 18 | H11 | 90 | 91 | - | - | This study |
| Cas9 | mRNA | Bovine | CI | 18 | ZFX | 67 | 100 | - | - | This study |
| Cas9 | RNP | Bovine | CI | 18 | ZFX | 81 | 88 | - | - | This study |

In bovine embryos, DNA replication occurs approximately 12-14 hours after fertilization^11^. When injecting at 18hpi, as is often done when using traditional *in-vitro* fertilization (IVF) protocols, most zygotes would be expected to have completed DNA replication^12^ and there would likely be more than two copies of each chromosome, thus more opportunities for multiple genomic edits to occur, resulting in mosaicism. Additionally, following cytoplasmic injection, the gRNA/Cas9 ribonucleoprotein (RNP) complex needs time to enter the nucleus, find its target and cleave the DNA. Furthermore, if injecting Cas9 mRNA, translation to Cas9 protein must also occur, further delaying the editing process, thus resulting in a higher rate of mosaicism. It has been suggested that injection of the CRISPR/Cas9 RNP prior to the S-phase of DNA replication could reduce mosaicism^1^.

One recent study with bovine embryos reported low rates (~30%) of mosaicism when introducing Cas9 RNA or protein into early stage zygotes (0 or 10hpi) prior to the S-phase of DNA replication^12^. In that study, allele identification was first made by Sanger sequencing of an amplicon of the targeted region, and then by clonal sequencing of 10 colonies derived from the PCR product per embryo. PCR and cloning-based approaches can identify that a range of alleles exist but cannot accurately quantitate the abundance of each allelic species. The authors went on to employ next generation sequencing on 20 embryos per group to characterize the alleles in non-mosaic embryos. The authors considered embryos that contained biallelic mutations resulting in a gene knockout to be non-mosaic, regardless of the proportion of alleles.

In the current study, we employed next generation sequencing to quantitate the abundance of each allele. The fact that we observed multiple alleles occurring in only a small percentage of reads (< 25%) in many samples analyzed in this study (Figure 2) suggests that editing continued in some subset of cells after the first cleavage division. Further, we considered an embryo containing more than one population of genetically distinct cells to be mosaic irrespective of whether the edit resulted in a missense or nonsense mutation. It is important to determine if founder animals are mosaic because mosaicism complicates the interpretation of the effect of a given genome alteration^5^, and subsequent breeding of mosaic founder animals to achieve non-mosaic animals can take years^13^. Additionally, mosaics do not fit easily into the proposed regulatory framework for genome edited food animals^14^.

Along with the level of mosaicism, one of the concerns raised with the generation of genome edited animals is the potential for off-target mutation events. Typically, online prediction tools are used to calculate the likelihood of off-target sites^15-17^. The top predicted sites can then be PCR amplified and the presence of a mutation determined by either next generation sequencing, TA cloning followed by Sanger sequencing, or mismatch cleavage assays followed by Sanger sequencing^18^. In this study, we used the targeted approach using online predictive tools to identify off-target sites rather than a genome-wide approach. Off-target cleavage can occur in the genome with three to five base pair mismatches in the PAM-distal sequence^15,19-21^. Cas9 specificity is determined by the seed region, or the 8 to 11-nt PAM-proximal sequence, making it the most vital part of the gRNA sequence^19,22^. In our gRNA design, we excluded all gRNAs with less than three mismatches across the off-target sequence. We determined this threshold based on previous studies showing reduced Cas9 activity in regions with at least three mismatches^23^.

In the 69 samples analyzed, there were two potential off-target mutations detected. One of these (H11) was in a region that had known annotated wild type 12bp deletions (rs876383581 and rs521367917) around the potential cut-site. Additionally, 0.51% of total reads contained a 3bp deletion 11bp downstream from the predicted off-target cut site for the ZFX gRNA target chr21: 28506796 (Supplemental Information, Table S7). This predicted site does not have any annotated variation. It is important to note that although this off-target location had three mismatches to the gRNA sequence, all three of the mismatches were located outside the seed region (8-11bp upstream of the PAM sequence). This guide was designed using off-target prediction software and the Btau 4.6.1 bovine reference genome^24^, which was the only *Bos taurus* reference genome available with the online tool at the time. When the off-target prediction software was re-run for the off-target analysis, the most recent reference genome available was UMD 3.1.1^24^. Using the new reference genome, this locus on chromosome 21 was identified as having the requisite three mismatches, but there were no mismatches in the seed region, as specified by our guide design criteria. More recently, an improved reference bovine genome ARS-UCD1.2 was published^25^. Using the online tool with the updated reference genome resulted in the same predicted off-target sites as UMD 3.1.1.

One of the stated concerns with off-target mutation events is that if they occur in functional regions, such as coding sequences or regulatory regions, they could potentially be detrimental to the health or development of the resulting animal. Neither of these two off-target deletions were in a region of annotated function. As there were approximately 20 individual blastocysts included in these analyses, these deletions may also have been the result of naturally occurring polymorphic variation. A detailed sequence analysis of 2,703 individuals from different breeds of cattle revealed a high level of genetic diversity including 84 million single-nucleotide polymorphisms (SNPs) and 2.5 million small insertion deletions^26^. Data like these are essential to put naturally occurring variation, like that seen at the H11 locus, in context. Various studies in humans^27,28^, monkeys^29^, and rodents^30,31^ suggest that the off-target frequency of Cas9-mediated mutagenesis does not differ from the de novo mutation rate.

Overall, we demonstrated efficient CRISPR/Cas9 genome editing across three different loci on three different chromosomes. We found that injecting zygotes with Cas9 protein results in a significantly higher mutation rate compared to Cas9 mRNA (82.2% vs 65.4%). In addition, zygotes injected with Cas9 protein displayed a significantly lower number of alleles compared to those injected with Cas9 mRNA (4.2 vs 5.2). Although off-target events did not appear to be an issue, the rate of mosaicism was still high, and further optimization needs to be done before this technique is feasible in a livestock production setting.

## MATERIALS AND METHODS

### Guide Construction

Guides sequences were designed using the online tools sgRNA Scorer 2.0^32,33^ and Cas-OFFinder^34^ and targeting the POLLED locus on chromosome 1, a safe harbor locus (H11) on chromosome 17 and in the 3’ UTR of the Zinc-finger X-linked (*ZFX*) gene (ZFX) on the X-chromosome. Guides were selected with no less than three mismatches in the guide sequence for off-target sites using the UMD3.1.1 bovine reference genome^24^, and at least one mismatch in the seed region (8-11bp upstream of the PAM sequence). Oligonucleotides were ordered from Eurofins USA (Louisville, KY) for the top four guides for construction of the gRNA and were used for *in vitro* transcription using the AmpliScribe T7-Flash Transcription kit (Lucigen, Palo Alto, CA) and purified using the MEGAclear Transcription Clean-Up kit (Thermo Fisher, Chicago, IL) as described by Vilarino et al^10^. Cleavage efficiency was tested using an *in vitro* cleavage assay by combining 60ng of PCR amplified product, 100ng of gRNA, 150ng of Cas9 protein (PNA Bio, Inc., Newbury Park, CA), 1μL of 10X BSA, 1μL of NEB Buffer 3.1 and water bringing the total volume to 10μL in a 0.2μL tube and incubating at 37°C for 1 hour. The incubated product was then run on a 2% agarose gel with 5μL of Sybr Gold at 100V for 1 hour and visualized using a ChemiDoc-ItTS2 Imager (UVP, LLC, Upland, CA).

### Embryo Production

Bovine ovaries were collected from a local processing plant and transported to the laboratory at 35-37°C in sterile saline. Cumulus-oocyte complexes (COCs) were aspirated from follicles and groups of 50 COCs were transferred to 4-well dishes containing 400μL of maturation media35. COCs were incubated for 21-24hr at 38.5°C in a humidified 5% CO_2_ incubator. Approximately 25 oocytes per drop were fertilized in 60μL drops of SOF-IVF^35^ with 1×10^6^ sperm per mL and incubated for 18hr at 38.5°C in a humidified 5% CO_2_ incubator. Presumptive zygotes were denuded by light vortex in SOF-HEPES medium^35^ for 5 min. 25 zygotes per drop were incubated in 50μL drops of KSOM culture media (Zenith Biotech, Glendale, CA, USA) at 38.5°C in a humidified atmosphere of 5% CO_2_, 5% O_2_, and 90% N_2_ for 7-8 days.

### Guide Testing

Mutation rate for each guide was determined by laser-assisted cytoplasmic injection^36^ of *in vitro* fertilized embryos with 6pL of a solution containing 67ng/μL of *in vitro* transcribed gRNA alongside 133ng/μL of Cas9 mRNA or 167ng/μL of Cas9 protein (PNA Bio, Inc., Newbury Park, CA) incubated at room temperature for 30 minutes prior to injection. Injected embryos were incubated for 7-8 days. Embryos that reached blastocyst stage were lysed in 10μL of Epicenter DNA extraction buffer (Lucigen, Palo Alto, CA) using a SimpliAmp Thermal Cycler (Applied Biosystems, Foster City, California) at 65°C for 6 minutes, 98°C for 2 minutes and held at 4°C. The target region was amplified by two rounds of the polymerase chain reaction (PCR) using primers developed using Primer3 (Supplementary Information, Table S1)^37,38^. The first round of PCR was performed on a SimpliAmp Thermal Cycler (Applied Biosystems, Foster City, California) with 10μL GoTAQ Green Master Mix (Promega Biosciences LLC, San Luis Obispo, CA), 0.4μL of each primer at 10mM and 9.2μL of DNA in lysis buffer for 5 min at 95°C, 35 cycles of 30 sec at 95°C, 30 sec at anneal temp (Supplementary Information, Table S1), and 30 sec at 72°C, followed by 5 min at 72°C. The second round of PCR was run with 10μL GoTAQ Green Master Mix (Promega Biosciences LLC, San Luis Obispo, CA), 4.2μL of water, 0.4μL of each primer at 10mM and 5μL of first round PCR for 3 min at 95°C, 35 cycles of 30 sec at 95°C, 30 sec at anneal temp (Supplementary Information, Table S1), and 30 sec at 72°C, followed by 5 min at 72°C. Products were visualized on a 1% agarose gel using a ChemiDoc-ItTS2 Imager (UVP, LLC, Upland, CA), purified using the QIAquick Gel Extraction Kit (Qiagen, Inc., Valencia, CA) and Sanger sequenced (GeneWiz, South Plainfield, NJ).

### Allelic Variation and Off-Target Analysis

Embryos that reached the blastocyst stage were lysed and underwent whole-genome amplification using the Repli-G Mini kit (Qiagen, Inc., Valencia, CA). To determine presumptive off-target sites, guide sequences were mapped against the bosTau8 bovine reference genome using the online tool Cas-OFFinder^34^. A total of 24 off-target sites were predicted using the online tool: eight off-target sites for the POLLED gRNA, eleven off-target sites for the H11 gRNA and five off-target sites for the ZFX gRNA (Supplementary Information, Table S7). Whole-genome amplified samples were used for PCR amplification of cut-sites and presumptive off-target sites using a dual round PCR approach described above to barcode each sample. Primers were designed to amplify each region using Primer3^37,38^ with a 15bp adapter sequence attached to the forward (AGATCTCTCGAGGTT) and reverse (GTAGTCGAATTCGTT) (Supplementary Information, S1). The second round of PCR amplified off the adapters adding an independent barcode for each sample to identify reads for pooled sequencing (Supplementary Information, Table S1).

PCR samples targeting the gRNA cut site underwent SMRTbell library preparation and were sequenced on a PacBio Sequel II sequencer by GENEWIZ, LLC (South Plainfield, NJ, USA). Consensus sequences were called, reads sorted by barcode and BAM converted to individual FASTQ files using SMRT Link v8.0.0.80529 (https://www.pacb.com/support/software-downloads/). Reads were aligned to each target site using BWA v0.7.16a^39^. SAM files were converted to BAM files, sorted and indexed using SAMtools v1.9^40^. Number and types of alleles were determined for each sample using CrispRVariants v1.12.0^41^.

Off-target PCR samples underwent library preparation using the Illumina TruSeq library kit and were sequenced (300bp paired-end) on an Illumina MiSeq Next Generation Sequencer by the DNA Technologies and Expression Analysis Cores at the UC Davis Genome Center. Paired-end reads were processed and overlapped to form high quality single-end reads using HTStream Overlapper v1.1.0 (https://github.com/ibest/HTStream). Processed reads were aligned to each target site using BWA v0.7.16a^39^. SAM files were converted to BAM files, sorted and indexed using SAMtools v1.9^40^. Insertions and deletions were called using CrispRVariants v1.12.0^41^.

### Statistical Analysis

Comparison between development for guide analysis and mutation rates were evaluated using a linear model and statistical significance was determined using a Chi-square test. To analyze the level of mosaicism, an ANOVA test was used to determine significance between number of alleles per sample and percent wild type when injecting alongside Cas9 mRNA or protein. Samples with only wild type alleles were removed from analysis. Differences were considered significant when P < 0.05.

## Acknowledgements

The authors would sincerely like to thank Ashley Young and Rebecca Ozeran for diligent collection and transport of ovaries from the processing plant to UC Davis. The library preparation and sequencing for off-target analysis were carried out at the DNA Technologies and Expression Analysis Cores at the UC Davis Genome Center, supported by NIH Shared Instrumentation Grant 1S10OD010786-01. The authors would like to thank the University of California Davis Bioinformatics Core Facility for assistance with data analysis. This project was supported by Biotechnology Risk Assessment Grant Program competitive grant no. 2015-33522-24106 from the U.S. Department of Agriculture, the Academic Federation Innovation Development Award at UC Davis, the Russell L. Rustici Rangeland and Cattle Research Endowment in the College of Agricultural and Environmental Science at UC Davis, the California Agricultural Experiment Station of the University of California, Davis, the Henry A. Jastro Research Fellowship in the College of Agricultural and Environmental Science at UC Davis and the National Institute for Food and Agriculture National Needs Graduate and Postgraduate Fellowship no. 2017-38420-26790 from the U.S. Department of Agriculture.

## Author’s contribution

SLH and JRO performed the experiments with additional input from PJR, ALV and JDM. SLH, JRO, JCL and AEY participated in sample processing and data analysis. SLH, JRO, ALV and JDM wrote the manuscript with suggestions from all the co-authors. All authors read and approved the final version.

## Data Availability

Raw sequence reads from PacBio Sequel II and Illumina MiSeq sequencing are available in the NCBI Sequence Read Archive as BioProject PRJNA623431 and SRA accession number SRR11850065. Individual results for the blastocyst development and mutation rate from each replicate (~ 30 embryos) of control and microinjected embryos are available in Supplementary Table S8.

## Supplementary Data

**Supplementary Table S1.**
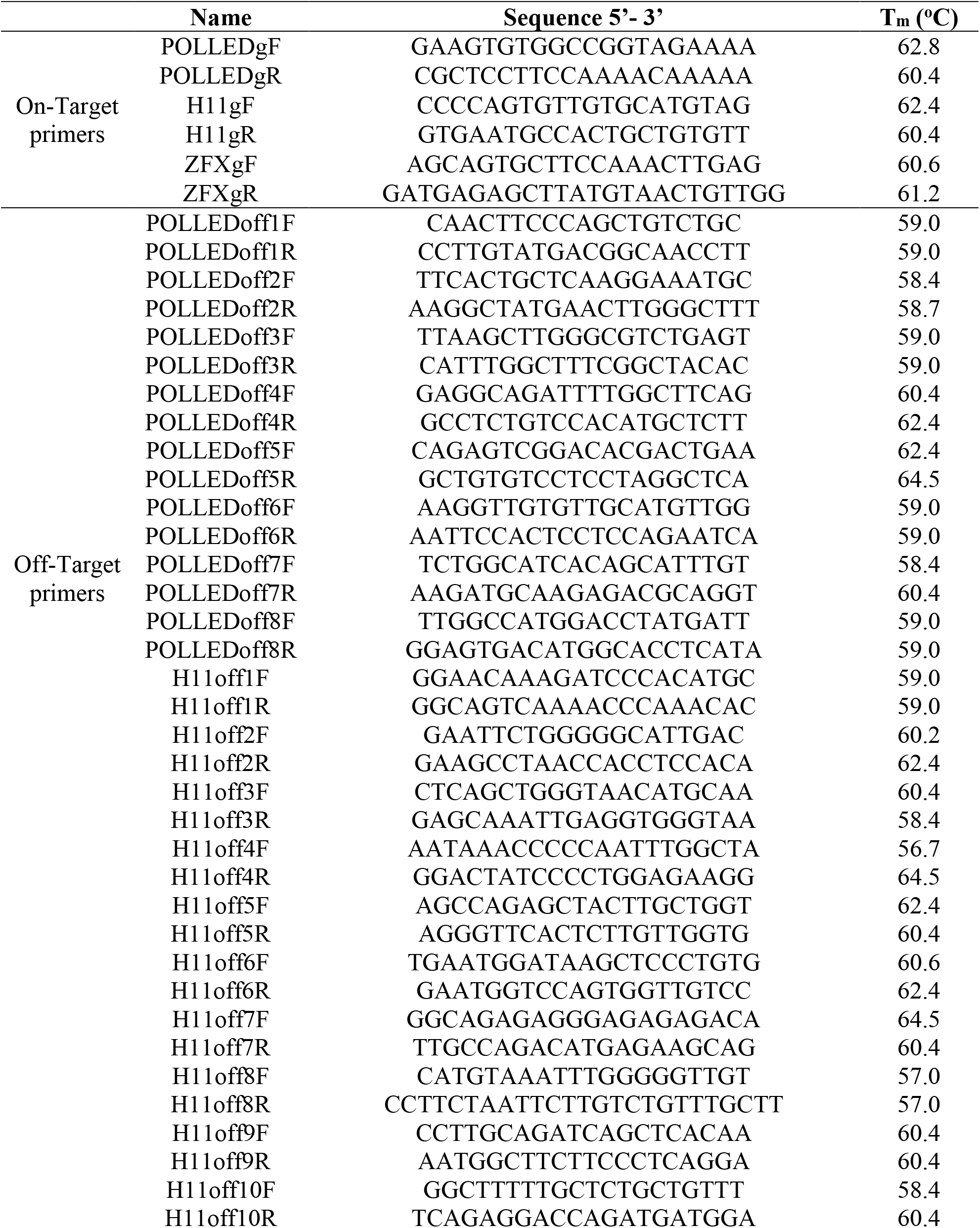
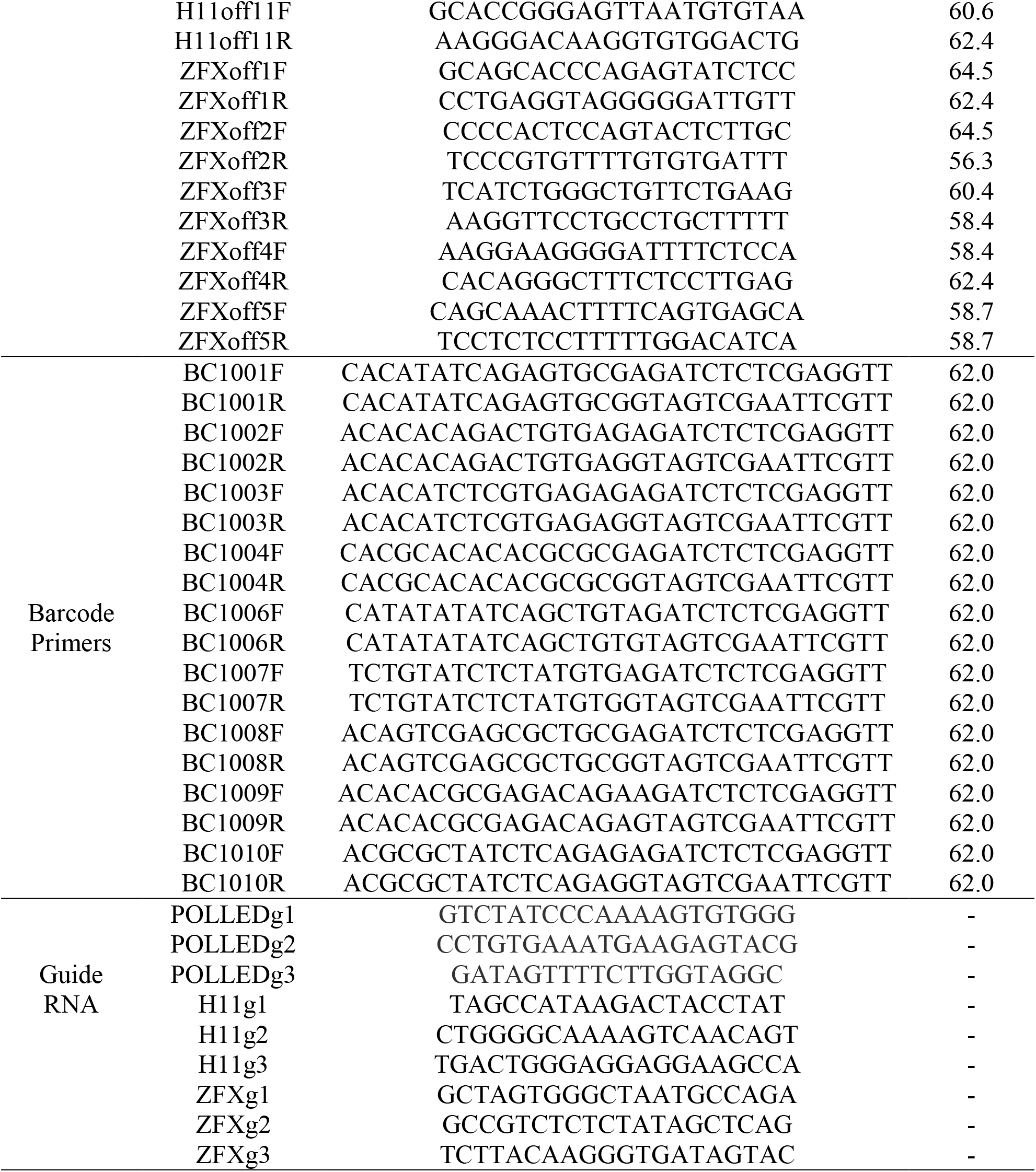
Sequence of primers used for PCR amplification of the POLLED, H11, or ZFX target regions, predicted off-target regions and gRNA sequences.

**Supplementary Table S2.** Mutation rate in embryos for each guide injected 18 hours post insemination alongside Cas9 protein analyzed using PCR and Sanger sequencing. Multiple guides were tested targeting each locus to obtain highest efficiency guide. Letters that differ in the same column are significantly different (P < 0.05). Each chromosome independently tested using a two-by-two χ^2^ test.

| Allele | gRNA | Injected Embryos | Total Blastocysts (%) | Blastocysts Analyzed | Mutation Rate (%) |
| --- | --- | --- | --- | --- | --- |
| POLLED | 1 | 47 | 15 (32<sup>a</sup>) | 13 | 0 (0)<sup>a</sup> |
|  | 2 | 75 | 14 (19<sup>b</sup>) | 13 | 10 (77)<sup>b</sup> |
|  | 3 | 90 | 25 (28<sup>a</sup>) | 25 | 2 (8)<sup>a</sup> |
| H11 | 1 | 65 | 12 (18<sup>b</sup>) | 12 | 10 (83)<sup>b</sup> |
|  | 2 | 45 | 13 (29<sup>a</sup>) | 13 | 5 (38)<sup>a</sup> |
|  | 3 | 47 | 10 (21<sup>b</sup>) | 10 | 6 (60)<sup>ab</sup> |
| ZFX | 1 | 75 | 22 (29<sup>a</sup>) | 19 | 1 (5)<sup>a</sup> |
|  | 2 | 86 | 22 (26<sup>a</sup>) | 21 | 5 (24)<sup>a</sup> |
|  | 3 | 104 | 18 (17<sup>b</sup>) | 18 | 14 (78)<sup>b</sup> |

**Supplementary Table S3.**
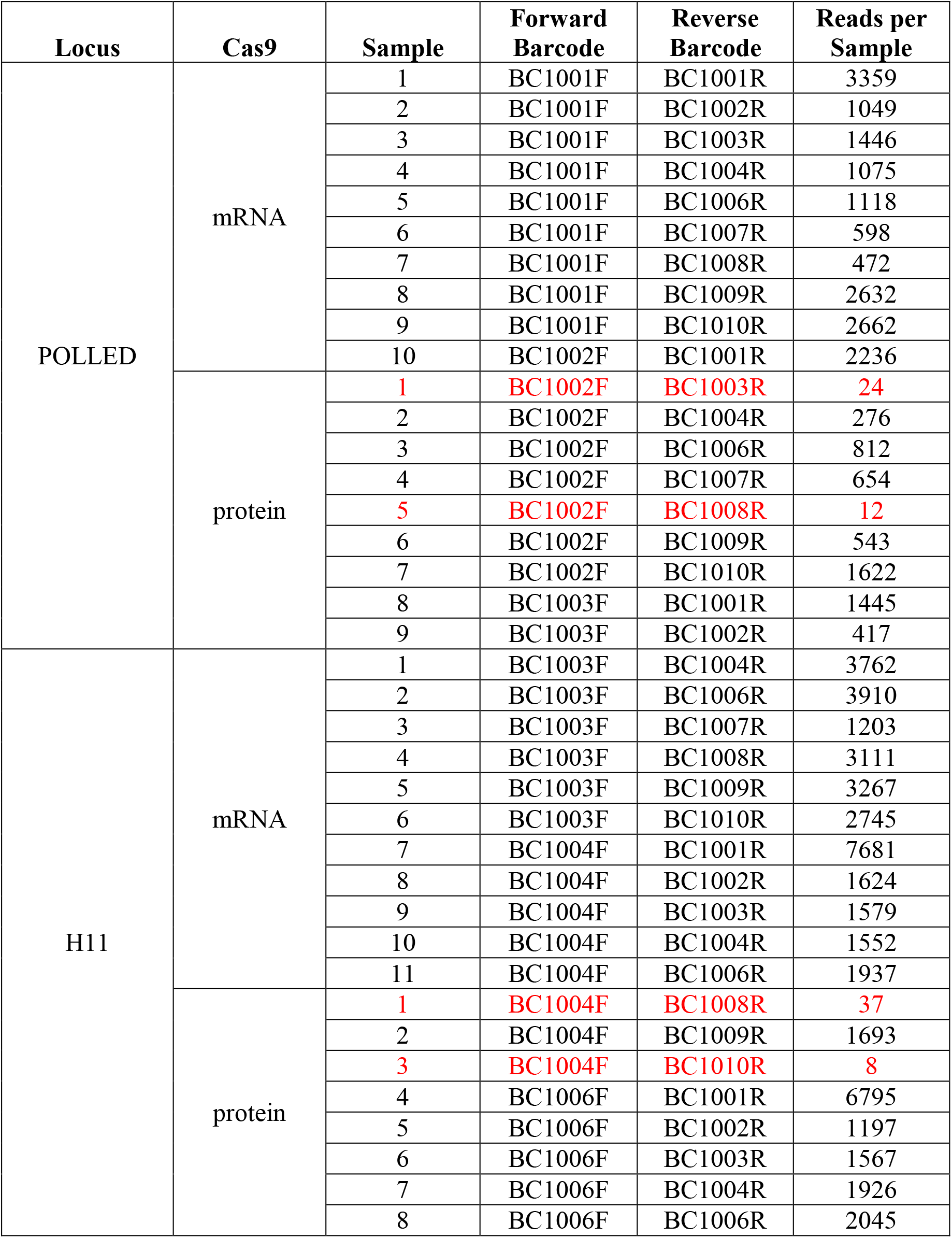
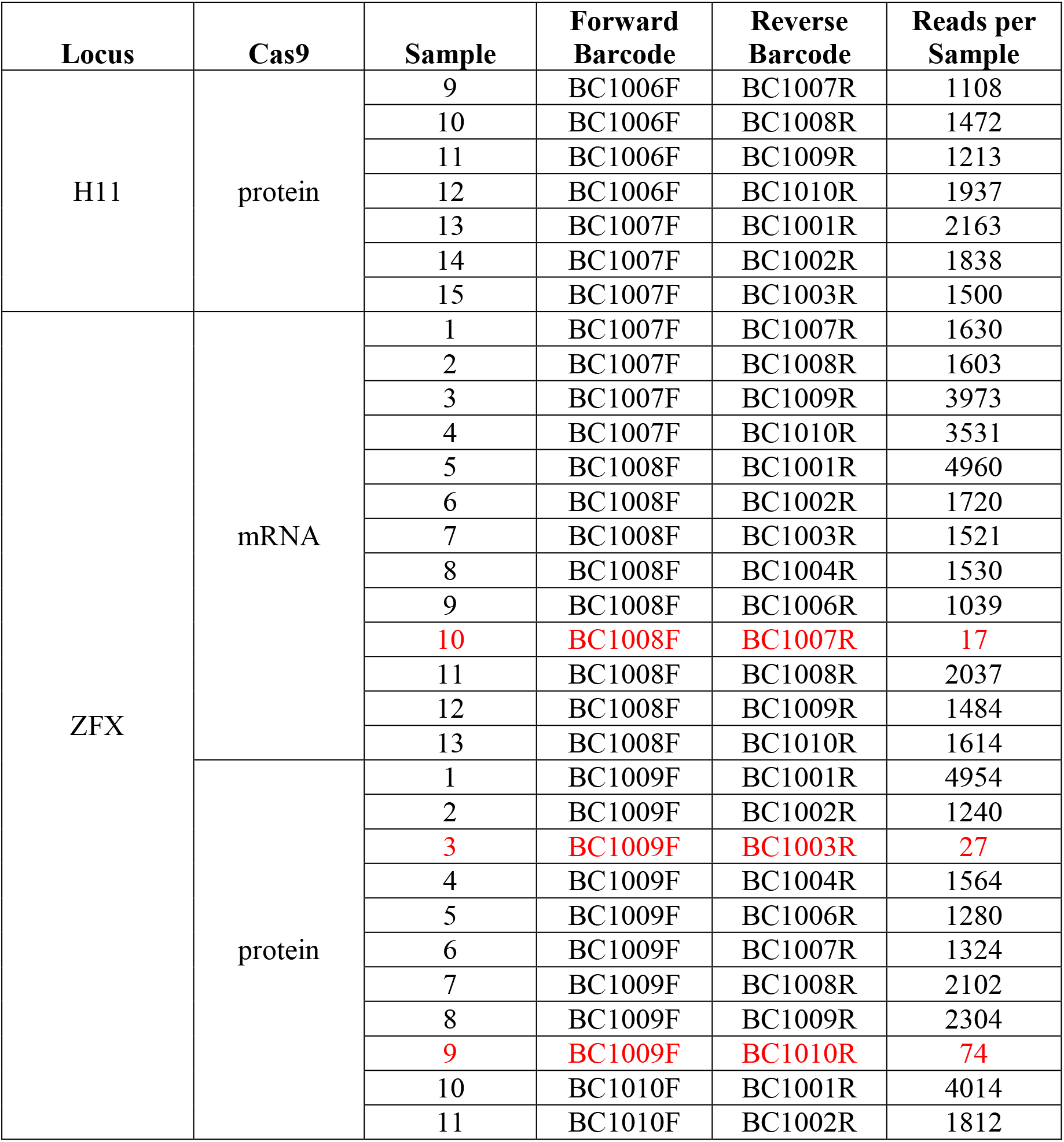
List of sequencing barcodes used for PacBio sequencing for embryos injected 18 hours post insemination with gRNAs targeting the POLLED, H11, and ZFX loci alongside Cas9 mRNA or protein and corresponding reads per sample following sorting by barcode. Red highlighted samples were removed from analysis due to insufficient read count.

**Supplementary Table S4.** Number of PacBio sequencing reads of PCR products from 69 blastocysts microinjected with Cas9 editing reagents targeting three loci (POLLED, H11, and ZFX) in the bovine genome, and the percentage of reads that were <700 bp read length, and additionally had a unique blastocyst sample identifying barcode.

| Filtered<br>By | Locus | Passed | Total | Percent |
| --- | --- | --- | --- | --- |
| Read length | Total | 171,545 | 236,518 | 72.5 |
| Barcode | POLLED | 22,416 | 26,460 | 84.7 |
| Barcode | H11 | 58,815 | 78,305 | 75.1 |
| Barcode | ZFX | 47,236 | 66,780 | 70.7 |

**Supplementary Table S5.** Prevalence of different allele types from PacBio sequencing of targeted PCR products < 700 bp from 69 blastocysts microinjected with Cas9 editing reagents targeting three loci (POLLED, H11, and ZFX) in the bovine genome. Types of mutations = location relative to the cut site (3bp upstream of the PAM sequence): type of deletion; D = deletion, I = insertion. “Other” mutations indicate those with reads too few to report.

| Locus | Total Number<br>of Reads | Wild Type<br>Alleles (%) | Type of Mutation |  |  |  |  |
| --- | --- | --- | --- | --- | --- | --- | --- |
|  |  |  | # of Reads (%) |  |  |  |  |
| POLLED | 26460 | 12719 (48) | **-16:7D** | **-14:11D** | **-13:4D** | **-10:1I** | **Other** |
|  |  |  | 1672 (6) | 1751 (7) | 6356 (24) | 2250 (19) | 1712 (6) |
| H11 | 78305 | 59302 (76) | **-14:11D** | **-13:6D** | **-12:3D** | **-10:1D** | **-8:1D** |
|  |  |  | 3246 (4) | 3813 (5) | 4091 (5) | 5061 (6) | 2792 (4) |
| ZFX | 66780 | 47939 (71) | **-10:14D** | **-4:9D** | **-1:3D** | **1:1I** | **2:1I** |
|  |  |  | 4222 (6) | 2998 (4) | 3198 (5) | 2194 (3) | 6532 (10) |

**Supplementary Table S6.** Number of alleles and percentage of each corresponding allele per sample detected at the cut-site of Cas9 mRNA or protein injected embryos. WT = percentage of reads that were wild type sequence. Alleles 1-5 are percent reads with each of the alleles found in the samples. Bold samples contained no wild type sequence. n/a = not applicable; genotypic sex was only determined for samples targeting the X chromosome.

| Locus | Cas9 | Sample | Sex | # of alleles | % of Reads for Each Allele |  |  |  |  |  |
| --- | --- | --- | --- | --- | --- | --- | --- | --- | --- | --- |
|  |  |  |  |  | WT | Allele 1 | Allele 2 | Allele 3 | Allele 4 | Allele 5 |
| POLLED | mRNA | 1 | n/a | 5 | 57 | 22 | 10 | 8 | 3 | - |
|  |  | 2 | n/a | 5 | 76 | 11 | 7 | 4 | 2 | - |
|  |  | 3 | n/a | 6 | 37 | 44 | 9 | 6 | 4 | 3 |
|  |  | 4 | n/a | 5 | 9 | 31 | 28 | 25 | 8 | - |
|  |  | 5 | n/a | 6 | 46 | 22 | 20 | 6 | 6 | 4 |
|  |  | 6 | n/a | 5 | 67 | 17 | 11 | 3 | 2 | - |
|  |  | 7 | n/a | 4 | 9 | 75 | 13 | 2 | - | - |
|  |  | 8 | n/a | 6 | 20 | 43 | 27 | 7 | 4 | 3 |
|  |  | 9 | n/a | 6 | 53 | 23 | 13 | 7 | 4 | 3 |
|  |  | 10 | n/a | 6 | 39 | 21 | 21 | 15 | 5 | 4 |
|  | protein | 1 | n/a | 4 | 10 | 43 | 28 | 19 | - | - |
|  |  | 2 | n/a | 3 | 13 | 63 | 24 | - | - | - |
|  |  | 3 | n/a | 3 | 7 | 87 | 6 | - | - | - |
|  |  | 4 | n/a | 3 | 10 | 76 | 14 | - | - | - |
|  |  | **5** | n/a | **2** | - | **81** | **19** | - | - | - |
|  |  | 6 | n/a | 5 | 20 | 37 | 25 | 16 | 2 | - |
|  |  | **7** | n/a | **1** | - | **100** | - | - | - | - |
| H11 | mRNA | 1 | n/a | 6 | 54 | 34 | 4 | 3 | 3 | 3 |
|  |  | 2 | n/a | 6 | 21 | 44 | 20 | 6 | 5 | 4 |
|  |  | 3 | n/a | 5 | 14 | 65 | 17 | 2 | 2 | - |
|  |  | 4 | n/a | 6 | 46 | 27 | 14 | 7 | 3 | 3 |
|  |  | 5 | n/a | 5 | 72 | 16 | 6 | 4 | 2 | - |
|  |  | 6 | n/a | 4 | 84 | 13 | 2 | 1 | - | - |
|  |  | 7 | n/a | 4 | 93 | 4 | 2 | 1 | - | - |
|  |  | 8 | n/a | 1 | 100 | - | - | - | - | - |
|  |  |  |  |  | WT | Allele 1 | Allele 2 | Allele 3 | Allele 4 | Allele 5 |
| H11 | mRNA | 9 | n/a | 1 | 100 | - | - | - | - | - |
|  |  | 10 | n/a | 1 | 100 | - | - | - | - | - |
|  |  | 11 | n/a | 1 | 100 | - | - | - | - | - |
|  | protein | 1 | n/a | 1 | 100 | - | - | - | - | - |
|  |  | 2 | n/a | 6 | 53 | 12 | 10 | 10 | 9 | 6 |
|  |  | 3 | n/a | 6 | 10 | 50 | 13 | 13 | 8 | 6 |
|  |  | 4 | n/a | 6 | 14 | 53 | 17 | 7 | 7 | 3 |
|  |  | 5 | n/a | 1 | 100 | - | - | - | - | - |
|  |  | 6 | n/a | 6 | 18 | 48 | 25 | 4 | 3 | 2 |
|  |  | 7 | n/a | 4 | 8 | 54 | 29 | 9 | - | - |
|  |  | 8 | n/a | 4 | 8 | 59 | 28 | 5 | - | - |
|  |  | **9** | n/a | **1** | - | **100** | - | - | - | - |
|  |  | 10 | n/a | 5 | 11 | 67 | 10 | 8 | 4 | - |
|  |  | 11 | n/a | 6 | 10 | 30 | 25 | 13 | 12 | 9 |
|  |  | 12 | n/a | 5 | 4 | 42 | 38 | 14 | 3 | - |
|  |  | 13 | n/a | 4 | 16 | 43 | 31 | 10 | - | - |
| ZFX | mRNA | 1 | female | 5 | 9 | 75 | 6 | 6 | 4 | - |
|  |  | 2 | male | 6 | 51 | 43 | 2 | 1 | 1 | 1 |
|  |  | 3 | female | 5 | 76 | 9 | 9 | 4 | 3 | - |
|  |  | 4 | female | 3 | 97 | 2 | 1 | - | - | - |
|  |  | 5 | male | 6 | 85 | 5 | 4 | 4 | 1 | 1 |
|  |  | 6 | female | 6 | 54 | 16 | 15 | 8 | 3 | 3 |
|  |  | 7 | male | 1 | 100 | - | - | - | - | - |
|  |  | 8 | female | 3 | 92 | 4 | 4 | - | - | - |
|  |  | 9 | female | 6 | 85 | 7 | 3 | 3 | 2 | 2 |
|  |  | 10 | female | 6 | 89 | 4 | 3 | 2 | 2 | 1 |
|  |  | 11 | female | 1 | 100 | - | - | - | - | - |
|  |  | 12 | male | 1 | 100 | - | - | - | - | - |
|  | protein | 1 | male | 5 | 65 | 12 | 8 | 7 | 7 | - |
|  |  |  |  |  | WT | Allele 1 | Allele 2 | Allele 3 | Allele 4 | Allele 5 |
| ZFX | protein | 2 | male | 6 | 54 | 37 | 3 | 2 | 2 | 2 |
|  |  | 3 | female | 1 | 100 | - | - | - | - | - |
|  |  | 4 | female | 4 | 42 | 44 | 8 | 6 | - | - |
|  |  | 5 | male | 6 | 26 | 26 | 23 | 18 | 5 | 2 |
|  |  | **6** | male | **1** | **-** | **100** | **-** | **-** | **-** | **-** |
|  |  | 7 | male | 3 | 67 | 30 | 3 | - | - | - |
|  |  | 8 | female | 6 | 20 | 36 | 20 | 16 | 6 | 3 |
|  |  | 9 | female | 4 | 15 | 45 | 36 | 4 | - | - |

**Supplementary Table S7.**
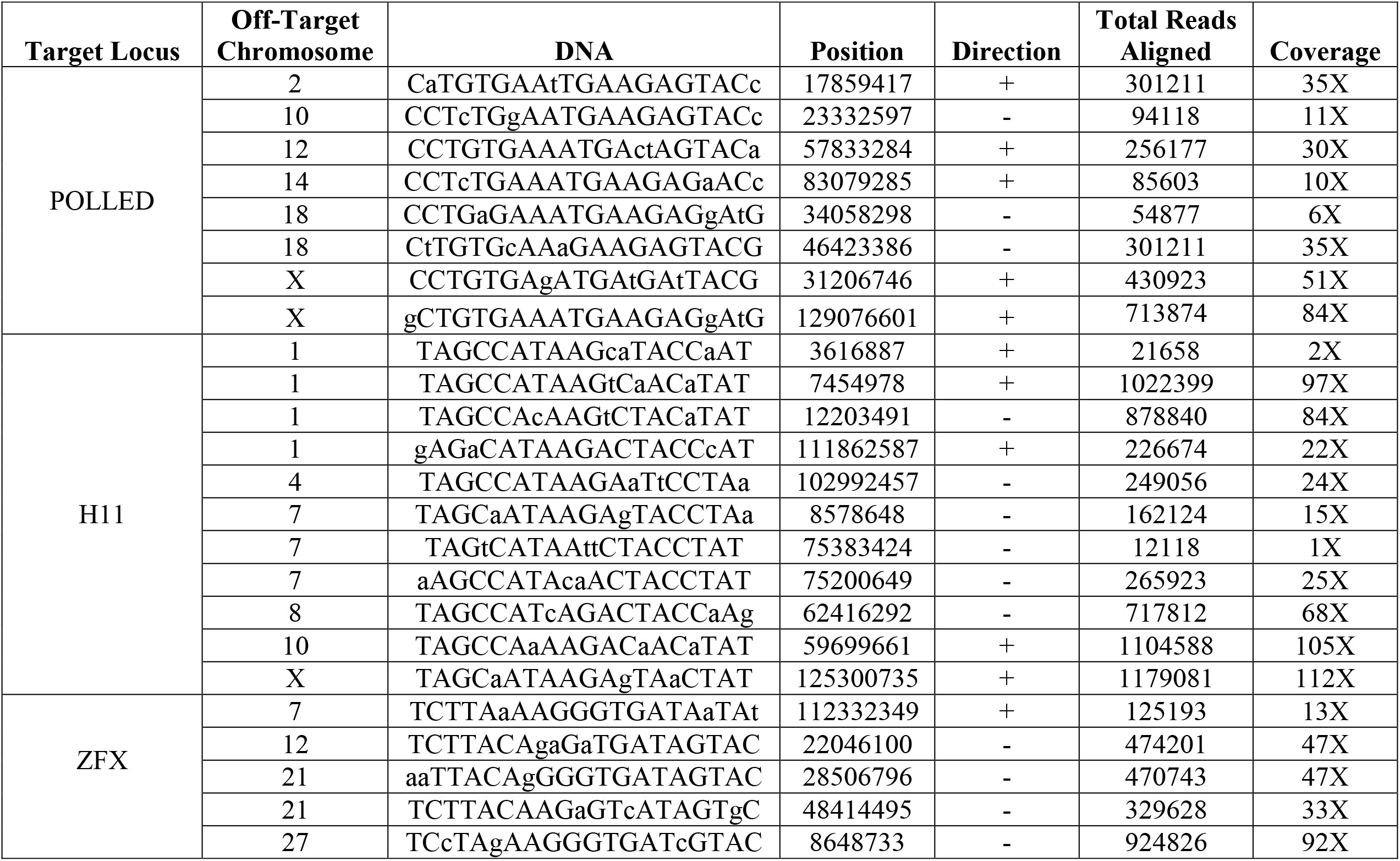
Predicted off-target sites for each of the three guides targeting the POLLED, H11, or ZFX locus. DNA = sequence of off-target site (lower case bases are mismatches). Position is relative to the start of the bosTau8 reference genome. Total reads aligned = number of reads mapped to the off-target sequence from overlapped MiSeq data. Coverage = reads per sample per target.

**Supplementary Table S8.** Results for development and mutation rate from each replicate of control embryos, and groups injected 18 hours post insemination with gRNAs targeting the POLLED, H11, and ZFX loci alongside Cas9 mRNA or protein

| Locus | Cas9 | # Embryos | # Blastocyst | % Blastocyst | # Blastocysts evaluated | # Blastocysts With mutation | % Mutation |
| --- | --- | --- | --- | --- | --- | --- | --- |
| POLLED | mRNA | 30 | 7 | 23.3 | 7 | 4 | 57.1 |
| POLLED | mRNA | 29 | 5 | 17.2 | 5 | 3 | 60 |
| POLLED | protein | 26 | 4 | 15.4 | 3 | 3 | 100 |
| POLLED | protein | 27 | 5 | 18.5 | 4 | 3 | 75 |
| POLLED | protein | 27 | 3 | 11.1 | 2 | 2 | 100 |
| POLLED | protein | 28 | 5 | 17.9 | 4 | 3 | 75 |
| POLLED | protein | 26 | 6 | 23.1 | 5 | 4 | 80 |
| POLLED | protein | 27 | 4 | 14.8 | 3 | 2 | 66.7 |
| POLLED | protein | 26 | 4 | 15.4 | 3 | 3 | 100 |
| POLLED | protein | 26 | 5 | 19.2 | 4 | 3 | 75 |
| POLLED | protein | 27 | 5 | 18.5 | 4 | 3 | 75 |
| POLLED | protein | 26 | 4 | 15.4 | 3 | 3 | 100 |
| H11 | mRNA | 26 | 3 | 11.5 | 3 | 2 | 66.7 |
| H11 | mRNA | 27 | 5 | 18.5 | 4 | 3 | 75 |
| H11 | mRNA | 26 | 3 | 11.5 | 3 | 2 | 66.7 |
| H11 | mRNA | 27 | 4 | 14.8 | 4 | 3 | 75 |
| H11 | mRNA | 25 | 3 | 12 | 3 | 2 | 66.7 |
| H11 | protein | 26 | 4 | 15.4 | 3 | 3 | 100 |
| H11 | protein | 26 | 5 | 19.2 | 4 | 3 | 75 |
| H11 | protein | 24 | 3 | 12.5 | 3 | 3 | 100 |
| H11 | protein | 26 | 4 | 15.4 | 3 | 3 | 100 |
| ZFX | mRNA | 26 | 4 | 15.4 | 4 | 3 | 75 |
| ZFX | mRNA | 26 | 4 | 15.4 | 4 | 2 | 50 |
| ZFX | mRNA | 27 | 5 | 18.5 | 5 | 3 | 60 |
| ZFX | mRNA | 26 | 5 | 19.2 | 5 | 3 | 60 |
| ZFX | mRNA | 26 | 4 | 15.4 | 4 | 3 | 75 |
| ZFX | mRNA | 27 | 4 | 14.8 | 4 | 2 | 50 |
| ZFX | mRNA | 26 | 3 | 11.5 | 3 | 2 | 66.7 |
| ZFX | mRNA | 25 | 5 | 20 | 5 | 4 | 80 |
| ZFX | mRNA | 26 | 3 | 11.5 | 3 | 1 | 33.3 |
| ZFX | mRNA | 25 | 4 | 16 | 4 | 3 | 75 |
| ZFX | mRNA | 26 | 5 | 19.2 | 5 | 4 | 80 |
| ZFX | mRNA | 26 | 4 | 15.4 | 4 | 2 | 50 |
| ZFX | protein | 26 | 4 | 15.4 | 4 | 3 | 75 |
| ZFX | protein | 26 | 4 | 15.4 | 4 | 3 | 75 |
| ZFX | protein | 27 | 5 | 18.5 | 5 | 4 | 80 |
| ZFX | protein | 26 | 5 | 19.2 | 5 | 5 | 100 |
| ZFX | protein | 26 | 4 | 15.4 | 4 | 3 | 75 |
| ZFX | protein | 24 | 4 | 16.7 | 4 | 3 | 75 |
| ZFX | protein | 26 | 3 | 11.5 | 3 | 3 | 100 |
| ZFX | protein | 25 | 5 | 20 | 5 | 4 | 80 |
| ZFX | protein | 26 | 3 | 11.5 | 3 | 2 | 66.7 |
| ZFX | protein | 25 | 4 | 16 | 4 | 3 | 75 |
| ZFX | protein | 26 | 5 | 19.2 | 5 | 4 | 80 |
| ZFX | protein | 26 | 4 | 15.4 | 4 | 4 | 100 |
| ZFX | protein | 26 | 4 | 15.4 | 4 | 3 | 75 |
| ZFX | protein | 25 | 5 | 20 | 5 | 4 | 80 |
| ZFX | protein | 25 | 4 | 16 | 4 | 3 | 75 |
| ZFX | protein | 26 | 4 | 15.4 | 4 | 4 | 100 |
| ZFX | protein | 24 | 3 | 12.5 | 3 | 2 | 66.7 |
| ZFX | protein | 25 | 4 | 16 | 4 | 3 | 75 |
| ZFX | protein | 26 | 4 | 15.4 | 4 | 3 | 75 |
| ZFX | protein | 26 | 3 | 11.5 | 3 | 2 | 66.7 |
| control | - | 30 | 9 | 30 | - | - | - |
| control | - | 30 | 8 | 26.7 | - | - | - |
| control | - | 30 | 7 | 23.3 | - | - | - |
| control | - | 30 | 6 | 20 | - | - | - |
| control | - | 30 | 13 | 43.3 | - | - | - |
| control | - | 30 | 8 | 26.7 | - | - | - |
| control | - | 30 | 10 | 33.3 | - | - | - |
| control | - | 29 | 9 | 31 | - | - | - |
| control | - | 30 | 8 | 26.7 | - | - | - |
| control | - | 30 | 13 | 43.3 | - | - | - |
| control | - | 30 | 8 | 26.7 | - | - | - |
| control | - | 30 | 9 | 30 | - | - | - |
| control | - | 30 | 7 | 23.3 | - | - | - |
| control | - | 30 | 8 | 26.7 | - | - | - |
| control | - | 27 | 9 | 33.3 | - | - | - |
| control | - | 30 | 10 | 33.3 | - | - | - |
| control | - | 30 | 8 | 26.7 | - | - | - |
| control | - | 30 | 5 | 16.7 | - | - | - |
| control | - | 29 | 7 | 24.1 | - | - | - |
| control | - | 30 | 6 | 20 | - | - | - |
| control | - | 30 | 8 | 26.7 | - | - | - |
| control | - | 29 | 9 | 31 | - | - | - |
| control | - | 30 | 8 | 26.7 | - | - | - |
| control | - | 30 | 14 | 46.7 | - | - | - |
| control | - | 30 | 8 | 26.7 | - | - | - |
| control | - | 28 | 13 | 46.4 | - | - | - |
| control | - | 30 | 8 | 26.7 | - | - | - |
| control | - | 30 | 7 | 23.3 | - | - | - |
| control | - | 29 | 8 | 27.6 | - | - | - |
| control | - | 30 | 9 | 30 | - | - | - |
| control | - | 30 | 12 | 40 | - | - | - |
| control | - | 30 | 11 | 36.7 | - | - | - |
| control | - | 30 | 9 | 30 | - | - | - |
| control | - | 28 | 9 | 32.1 | - | - | - |
| control | - | 30 | 10 | 33.3 | - | - | - |
| control | - | 30 | 13 | 43.3 | - | - | - |
| control | - | 30 | 7 | 23.3 | - | - | - |
| control | - | 26 | 8 | 30.8 | - | - | - |
| control | - | 27 | 9 | 33.3 | - | - | - |
| control | - | 30 | 12 | 40 | - | - | - |
| control | - | 29 | 11 | 37.9 | - | - | - |
| control | - | 30 | 10 | 33.3 | - | - | - |

**Supplementary Figure S1.**
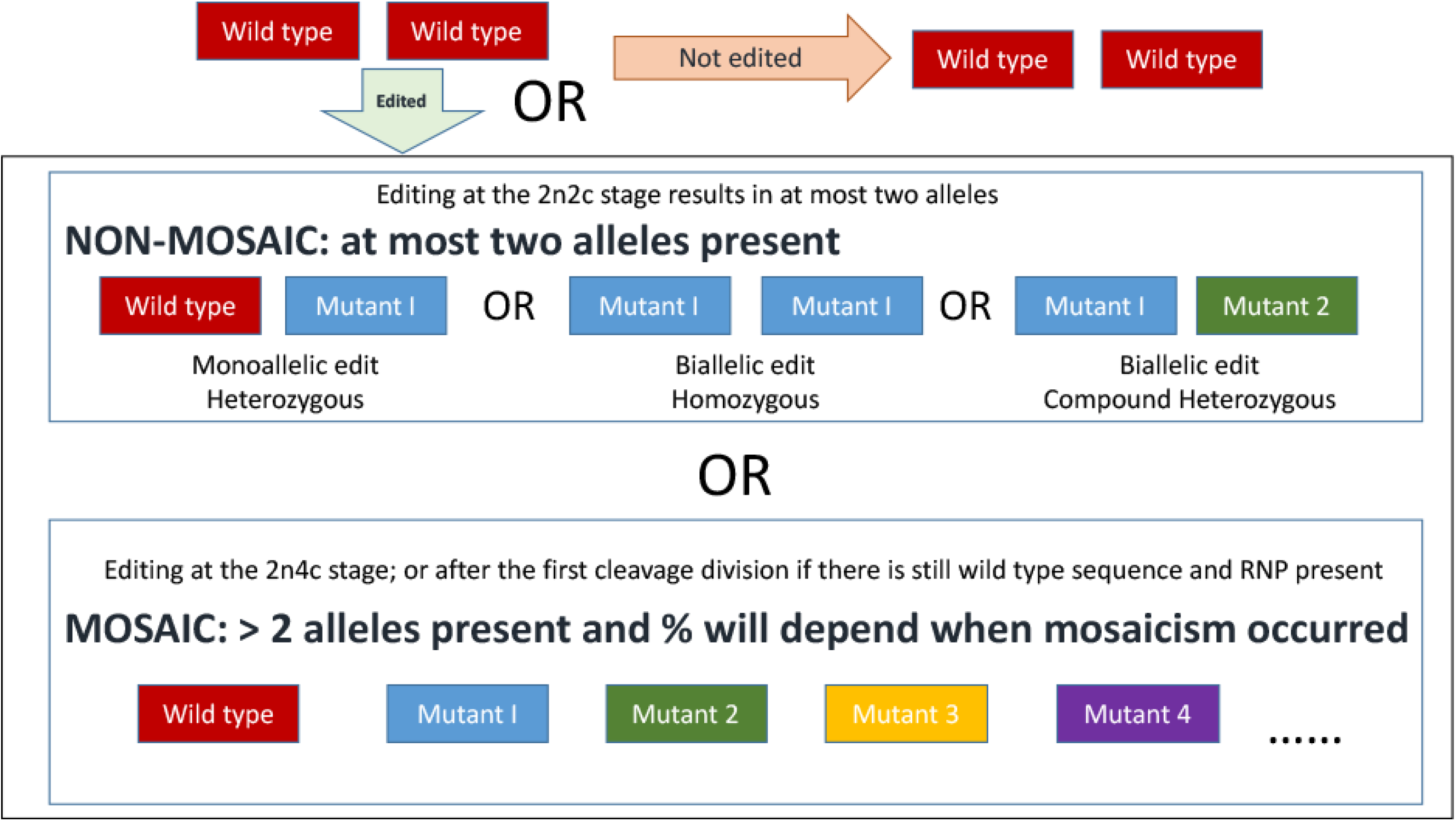
Schematic representation of possible outcomes from CRISPR-mediated mutation by cytoplasmic injection of an *in vitro* fertilized embryo 18 hours post insemination. 2n = number of homologous chromosomes, i.e. diploid. 2c/4c = number of copies of chromosomes either before DNA replication or after

